# The “Podcast” ECoG dataset for modeling neural activity during natural language comprehension

**DOI:** 10.1101/2025.02.14.638352

**Authors:** Zaid Zada, Samuel A. Nastase, Bobbi Aubrey, Itamar Jalon, Sebastian Michelmann, Haocheng Wang, Liat Hasenfratz, Werner Doyle, Daniel Friedman, Patricia Dugan, Lucia Melloni, Sasha Devore, Adeen Flinker, Orrin Devinsky, Ariel Goldstein, Uri Hasson

## Abstract

Naturalistic electrocorticography (ECoG) data are a rare but essential resource for studying the brain’s linguistic capabilities. ECoG offers a high temporal resolution suitable for investigating processes at multiple temporal timescales and frequency bands. It also provides broad spatial coverage, often along critical language areas. Here, we share a dataset of nine ECoG participants with 1,330 electrodes listening to a 30-minute audio podcast. The richness of this naturalistic stimulus can be used for various research endeavors, from auditory perception to semantic integration. In addition to the neural data, we extract linguistic features of the stimulus ranging from phonetic information to large language model word embeddings. We use these linguistic features in encoding models that relate stimulus properties to neural activity. Finally, we provide detailed tutorials for preprocessing raw data, extracting stimulus features, and running encoding analyses that can serve as a pedagogical resource or a springboard for new research.

## Background & Summary

We introduce the “Podcast” electrocorticography (ECoG) dataset for modeling neural activity supporting natural narrative comprehension. This dataset combines the exceptional spatiotemporal resolution of human intracranial electrophysiology with a naturalistic experimental paradigm for language comprehension. In addition to the raw data, we provide a minimally preprocessed version in the high-gamma spectral band to showcase a simple pipeline and to make it easier to use. Furthermore, we include the auditory stimuli, an aligned word-level transcript, and linguistic features ranging from low-level acoustic properties to large language model (LLM) embeddings. We also include tutorials replicating previous findings and serve as a pedagogical resource and a springboard for new research. The dataset comprises 9 participants with 1,330 electrodes, including grid, depth, and strip electrodes. The participants listened to a 30-minute story with over 5,000 words. By using a natural story with high-fidelity, invasive neural recordings, this dataset offers a unique opportunity to investigate language comprehension.

Language has a rich history of study in many scientific fields. Historically, research on the neural basis of language used highly controlled experimental stimuli to target particular linguistic phenomena, e.g., isolated words and sentences varying along a specific linguistic feature (Friederici, 2011; Price, 2012). The past decade has seen a push for a more ecological, holistic account of the neural basis of language processing (Hamilton & Huth, 2020; Hasson & Honey, 2012; Nastase et al., 2020). Yet naturalistic neural data is more challenging to model and understand than, for example, contrast-based experimental paradigms. Researchers have used data-driven methods like intersubject correlation (ISC) to isolate stimulus-driven processing in naturalistic contexts, sometimes under different conditions (Hasson et al., 2004; Nastase et al., 2019). However, data-driven methods like ISC do not allow us to test various models for the neural computations driving language processing (Goldstein et al., 2022; Yarkoni & Westfall, 2017; Zada et al., 2024).

Linear regression encoding models serve as a model-based analysis framework that maps linguistic and stimulus features to neural data (Holdgraf et al., 2017; Naselaris et al., 2011). These models learn a direct mapping from stimuli properties (e.g., word embeddings) to neural activity. With hundreds of regressors, regularization is often employed through PCA or a ridge penalty (Nunez-Elizalde et al., 2019). To evaluate their performance, encoding models are trained on a subset of data, then correlate model-based predictions with actual neural data for a held-out test set of data. If the prediction accuracy of the held-out test is high, we conclude that the neural activity encodes stimulus features represented by the model features. The encoding framework allows us to compare different stimulus properties, where different stimulus features act as hypotheses about the neural activity’s underlying function or representation.

With the current dataset, we offer five distinct models for processing the podcast data to evaluate on the ECoG dataset: 1) Low-level sensory features expressed as 80 mel-spectral power bins derived from the raw audio waveform. 2) Phoneme-level speech units are defined by decomposing each word into its constituent phonemes and represented using a 44-dimensional binary vector indicating the presence of a phoneme. In addition, we group consonants into the manner of articulation (e.g., nasal, plosive) and place of articulation (e.g., dental, palatal), based on de Heer et al., (2017). 3) Syntactic properties constituting 50 part-of-speech tags (e.g., nouns, verbs, adjectives, etc.), and 45 syntactic dependencies based on sentence parse trees codified as binary vectors. 4) Non-contextual word embeddings from pre-trained language models (e.g., GloVe (Pennington et al., 2014)), representing each word as a 300-dimensional vector representation learned from large corpora of natural text. Notably, each word is assigned the same vector regardless of its context. 5) Contextual word embeddings from a large language model (LLM) where each word vector is influenced by all preceding words within its contextual window. For example, we used GPT-2 XL (Radford et al., 2019), where word embeddings are 1,600-dimensional vectors. This tutorial demonstrates how to use these five models within an encoding comparison framework to predict neural activity, testing the effectiveness of competing models in capturing neural processes during natural language processing across different electrodes and brain areas.

The encoding-based model comparison showed that contextual word embeddings extracted from GPT-2 XL accounted for most of the variance across nearly all the electrodes tested in this dataset. These findings are aligned with recent work from our lab and others that find alignment between the neural activity in the human brain and the internal activity in LLMs during the processing of natural language (Hasson et al., 2020; Richards et al., 2019). For example, researchers used encoding models to show the brain’s predictive (Goldstein et al., 2022; Schrimpf et al., 2021) and hierarchical linguistic processing (Caucheteux et al., 2023; Heilbron et al., 2022); the shared underlying geometry (Bhattacharjee et al., 2024; Goldstein et al., 2024); the similarity between reading and listening (Deniz et al., 2019); the role of speech in language comprehension (Goldstein et al., 2023); and production-comprehension coupling between speakers (Zada et al., 2024).

The following describes the “Podcast” ECoG dataset, including data acquisition and preprocessing details. Next, we provide two quality control analyses to ensure the neural activity is precisely aligned with the onset of words in the podcast stimulus.

Finally, we use electrode-wise encoding analysis to test several linguistic features against the neural data.

## Methods

### Participants

Nine participants underwent clinical intracranial monitoring in the New York University School of Medicine’s Comprehensive Epilepsy Center. Participants consented to be included in our study following approval by the New York University Langone Medical Center’s Institutional Review Board. Clinicians determined electrode location and type per participant, based solely on the clinical need of the participant—without regard to any research study. Several standard behavioral tests were conducted to assess the patient’s memory and linguistic abilities. These include the Verbal Comprehension Index (VCI), Perceptual Organization Index (POI), Processing Speed Index (PSI), and Working Memory Index (WMI). Participant data is detailed in Table 1.

**Table 1.** Participant electrode, demographic, and behavioral information. Acronyms: left hemisphere (LH); right hemisphere (RH); verbal comprehension index (VCI); perceptual organization index (POI); processing speed index (PSI); and working memory index (WMI); ambidextrous (ambi.).

| subject |  | sub-01 | sub-02 | sub-03 | sub-04 | sub-05 | sub-06 | sub-07 | sub-08 | sub-09 |
| --- | --- | --- | --- | --- | --- | --- | --- | --- | --- | --- |
| electrode coverage | total | 103 | 95 | 264 | 155 | 160 | 171 | 119 | 75 | 188 |
|  | LH | n/a | 95 | 213 | 77 | 159 | 171 | 119 | 75 | 188 |
|  | RH | 103 | n/a | 51 | 78 | 1 | n/a | n/a | n/a | n/a |
|  | Grid | 63 | n/a | 127 | n/a | 110 | 127 | 63 | 47 | 124 |
|  | Strip | 24 | 43 | 65 | 79 | 8 | 8 | 32 | 8 | 40 |
|  | Depth | 16 | 52 | 72 | 76 | 42 | 36 | 24 | 20 | 24 |
| personal | sex | M | F | F | M | M | F | F | M | M |
|  | age | 28 | 40 | 48 | 25 | 24 | 34 | 28 | 42 | 24 |
|  | hand | R | R | R | R | ambi.<br>L>R | R | R | R | R |
| behavioral tests | VCI | 134 | 63 | 107 | n/a | 96 | n/a | 100 | 87 | 145 |
|  | POI | 127 | 79 | 104 | n/a | 79 | n/a | 109 | 123 | 96 |
|  | PSI | 124 | 84 | 111 | n/a | 81 | n/a | 92 | 97 | 86 |
|  | WMI | 119 | 89 | 114 | n/a | 86 | n/a | 86 | 89 | 95 |

Electrodes were made of platinum-iridium and were embedded in silastic sheets, with 2.3 mm diameter contacts (Ad-Tech Medical Instrument). Electrodes were grouped into three types: grid arrays, linear strips, or depth electrodes. Grid arrays consisted of 8 × 8 contacts with 5 or 10 mm center-to-center spacing. Three participants (sub-03, sub-06, sub-09) consented to a hybrid grid: a standard electrode grid augmented with additional electrodes between the standard contacts (i.e., a U.S. Food and Drug Administration-approved hybrid clinical research grid). In addition to the 64 standard contacts (2 mm diameter), the hybrid grid interleaved 64 additional contacts with 1 mm in diameter (these electrodes are labeled as the type “EG” in the dataset).

### Electrode localization

Each participant underwent a pre-surgical and post-surgical T1-weighted MRI scan. These two images were aligned following published methods (Yang et al., 2012). All accompanying anatomical scans are anonymized with *pydeface* (Gulban et al., 2022). Electrodes were then localized on the cortical surface from a co-registered MRI or computed tomography scan. Montreal Neurological Institute (MNI) electrode coordinates were obtained by nonlinearly registering the skull-stripped T1 images to the MNI152 template. Not all electrodes were on the brain; for example, some electrodes may be on the skull. We identified 80 electrodes that were not localized and were marked accordingly in the dataset.

### ECoG data acquisition

Electrodes were connected to one of two amplifier types: a NicoletOne C64 clinical amplifier (Natus Neurologics) that is band-pass filtered from 0.16–250 Hz, and digitized at 512 Hz; or a NeuroWorks Quantum Amplifier that is high-pass filtered at 0.01 Hz and recorded at 2,048 Hz. All signals were referenced to a two-contact subdural strip facing toward the skull near the craniotomy site.

### Stimuli

The stimulus presented to participants was a segment from the podcast “This American Life” entitled “So a Monkey and a Horse Walk Into a Bar: Act One, Monkey in the Middle” released on November 10, 2017. The original audio and transcript are freely available online (https://www.thisamericanlife.org/631/transcript).

### ECoG preprocessing

Raw electrode data underwent the following preprocessing pipeline: removing bad electrodes, downsampling, despiking and interpolating high amplitude spikes, common average re-referencing, and notch filtering. First, we visualized the power spectrum density of each electrode per subject. From this, we were able to annotate unusual electrodes that did not conform to the expected 1/f pattern, had a consistent oscillatory pattern, or showed other unusual artifacts. We found 31 such electrodes and marked them as “bad” (identified in the accompanying metadata). The source of these artifacts may be due to several factors, including excessive noise, epileptic activity, no noise, or poor contact. For data acquired with a sampling rate greater than 512 Hz, we downsampled to 512 Hz to match the sampling rate across subjects. We then applied a despiking and interpolation procedure to remove time points that exceeded four quartiles above the median of the signal and refill it using *pchip* interpolation. For re-referencing, we subtracted the mean signal across all electrodes per subject from each of their individual electrode time series. Finally, we used notch filters at 60, 120, 180, and 240 Hz to remove power line noise. This pipeline produces a “cleaned” version of the raw signal.

Linguistic processing may span multiple frequency bands and rely on cross-frequency coupling (e.g., Martin, 2020; Murphy, 2024). For the sake of simplicity, we focus on high-gamma band power as an index of local, stimulus-driven neuronal activity (Haufe et al., 2018; Manning et al., 2009; Mukamel et al., 2005). We extracted the high-gamma band by applying a Butterworth band-pass infinite impulse response (IIR) filter at 70–200 Hz. We extract the broadband power by computing the envelope of the Hilbert transform.

### Data Records

The dataset is freely available on OpenNeuro: https://openneuro.org/datasets/ds005574 (Markiewicz et al., 2021). Tutorials for usage are available at: https://hassonlab.github.io/podcast-ecog-tutorials. We followed the BIDS-iEEG (Gorgolewski et al., 2016; Holdgraf et al., 2019) standard for file structures and naming conventions. The main directory contains the raw ECoG data for each subject in EDF format under the *ieeg* datatype directory. In addition, channel information and electrode localization for MNI and T1 spaces are also located in the same folder, following BIDS guidelines. Notably, the channels tsv file contained annotation for each electrode in the EDF that denoted whether it’s a “good” or “bad” channel. Bad channels are those that are rejected either due to “no localization” or noisy power spectrum density (see “ECoG preprocessing” in the “Methods” section). Each subject also has a de-faced T1 anatomical MRI scan under their respective *anat* datatype directory.

The *derivatives* directory contains the preprocessed ECoG data in MNE .*fif* format within the “ecogprep” subdirectory (inspired by fMRIPrep; (Esteban et al., 2019)). There are two versions of the preprocessed data, one is unfiltered and the other is filtered to the high-gamma broadband power. In addition, the “ecogqc” subdirectory contains the output of a quality-check program in HTML that contains an interactive viewer for electrodes and the power spectrum density plots (inspired by MRIQC; (Esteban et al., 2017)). These were the primary sources used for determining the quality of an electrode’s signal.

The top-level *stimuli* directory contains the original audio presented to participants in .*wav* format, along with a time-stamped transcript in .*csv* format. Extracted data from each of the five feature spaces is placed in a subdirectory of *stimuli* denoting its type. For example, the *stimuli/spectral/* directory contains spectrogram features of the audio waveform. Each feature directory contains at least two files: *features*.*hdf5* and *transcript*.*tsv*. The *transcript* file is a modified version of the original transcript (under *stimuli*) where a particular feature type may remove or add additional rows or columns (e.g., for tokenization). The *features* file contains the numerical vectors of the feature space as a matrix, where the rows correspond one-to-one to the rows in the *transcript* file. The number of columns varies depending on the dimensionality of the feature space. In some cases, such as large language models, this file may include the activations of all layers in separate HDF5 groups in the same file. Examples of loading and manipulating these files can be found in the accompanying code and tutorials.

## Technical Validation

### Cross-correlation between electrode activity and auditory stimulus

The first analysis tested electrodes that track the acoustic properties of the stimulus, roughly aligning the neural activity with the audio. We cross-correlated the stimulus audio envelope with each electrode’s high-gamma broadband power. This analysis was used in Honey et al., (2012) to find electrodes that tracked the acoustic properties of the stimulus. Similar to their results, we also found a gradient of audio correlations that peaks in electrodes closest to the early auditory cortex and decrements along the anterior-posterior axis (Figure 2A). We also visualized the average cross-correlation of electrodes in the superior temporal cortex in a four-second window. This validation analysis reveals that brain activity lags just behind the auditory stimulus, demonstrating that the ECoG recordings are closely aligned with the stimulus.

### Electrode-wise evoked response to word onsets

We used an event-related potential (ERP) analysis as a complementary method to confirm the transcript word timings and validate the neural signal. We used the timed transcript to epoch the neural data (Figure 1D) and then averaged the neural signal in a four-second window around word onset. As expected, electrodes that are sensitive to word occurrence increased their activity from baseline (Figure 2B). These electrodes were localized in the vicinity of the early auditory cortex in the superior temporal cortex. The average evoked response across electrodes in the superior temporal cortex exhibited a sharp increase in activity around word onset.

**Figure 1.**
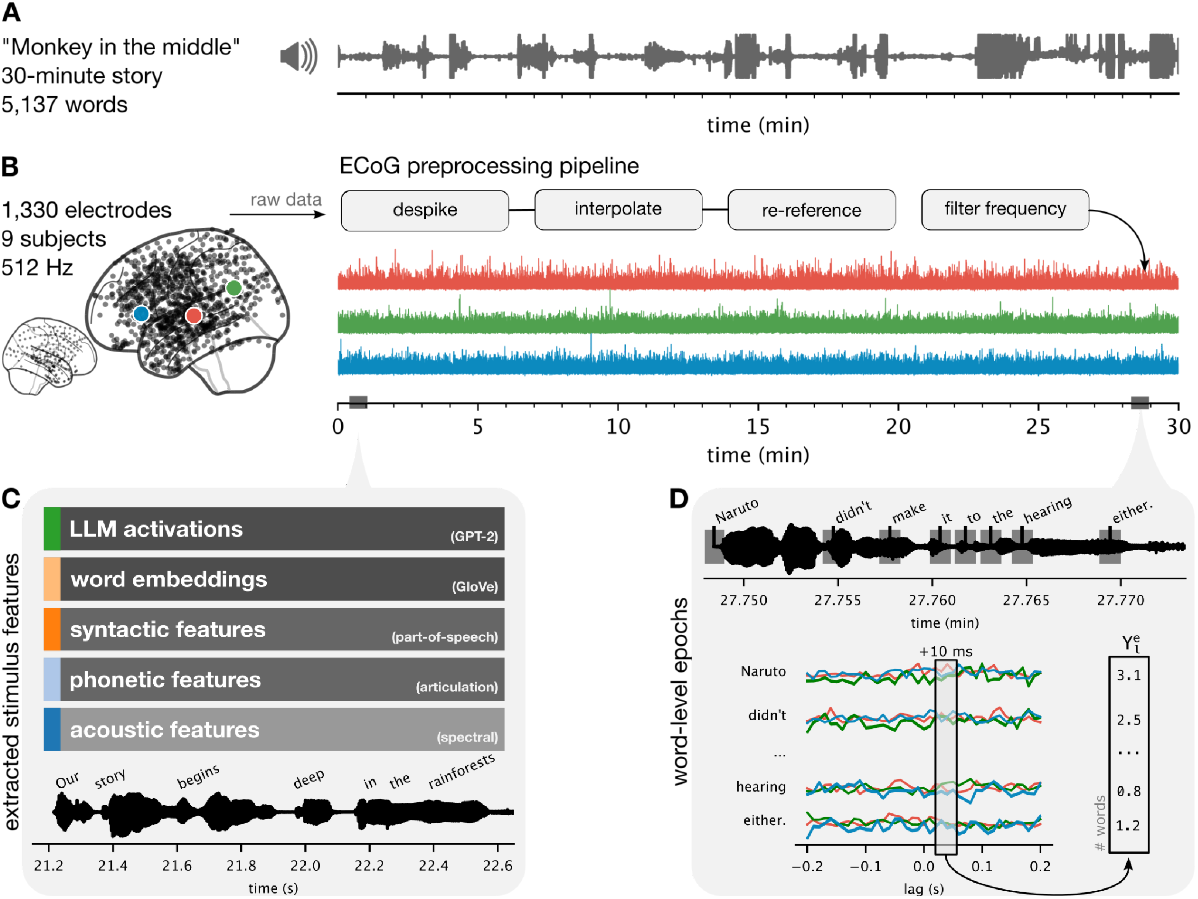
Experiment setup and dataset components. (**A**) A 30-minute audio story (podcast) was played to (**B**) nine participants while undergoing electrocorticographic monitoring for epilepsy. We implement a simple preprocessing pipeline to extract the high-gamma band power per electrode. (**C**) We manually transcribed the story and timestamped words at high temporal resolution. We extract five linguistic features from the audio waveform and transcript. (**D**) Some analyses depend on word-level epochs, where a window relative to word onset at each word is then stacked across words. This forms a matrix of words by the number of lags for each electrode. In electrode-wise encoding analysis, we use linear regression to predict the neural activity at a particular lag across words, separately for each electrode, depicted as a *Y* vector in (**D**), from the stimulus features defined in (**C**).

**Figure 2.**
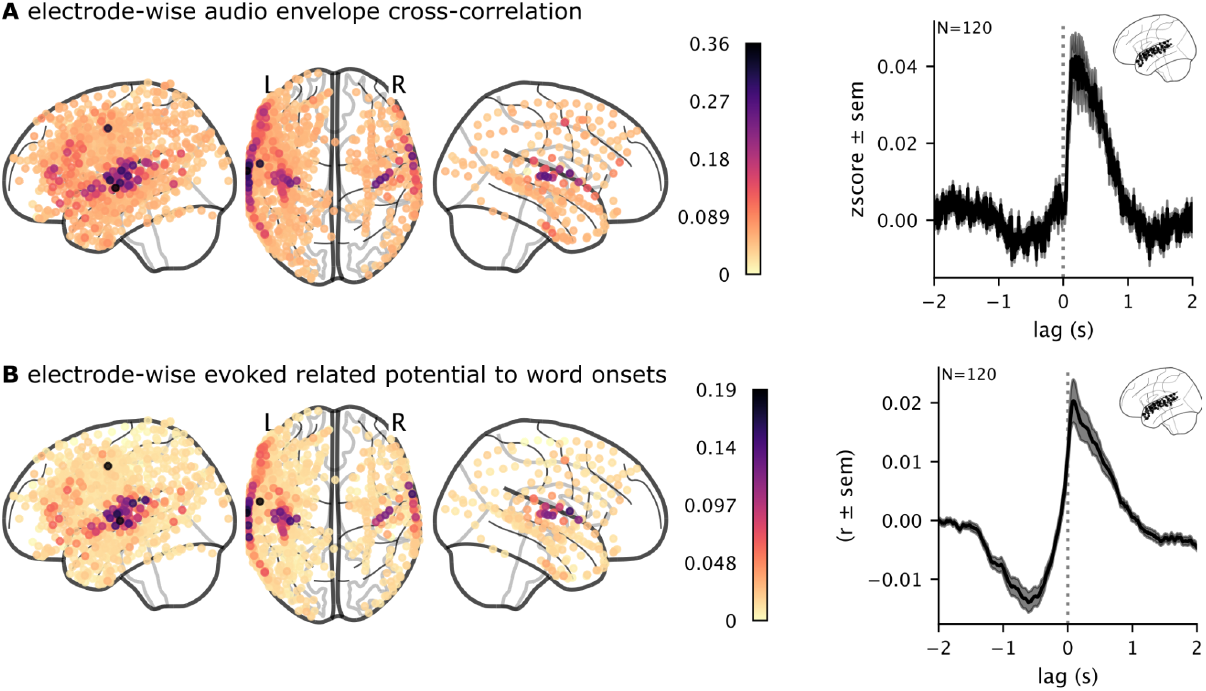
Validation of brain activity responses to auditory stimuli. (**A**) We cross-correlated the high-gamma band ECoG signal with the envelope of the audio waveform to validate the alignment between the audio and the brain data, especially in the early auditory cortex. (**B**) For each electrode, we computed the average neural responses across words to verify that electrodes in the early auditory cortex exhibit increased word-related activity.

### Electrode-wise encoding models for multiple feature spaces

We used electrode-wise encoding models to measure the neural responses to specific features of the podcast stimulus. We extracted five sets of linguistic features from the stimulus (Figure 1C). First, we computed a spectrogram from the audio waveform based on 80 log mel-spectral bins. We then epoched the spectrogram according to word onsets from lag 0 ms to 200 ms and downsampled each epoch to a sampling rate of 10 Hz. Thus, each word was represented by 10 frames of 80 bins, flattened into an 800-dimensional vector. Second, we defined a phoneme-based word representation using a word-to-phoneme dictionary to create a 40-dimensional binary vector of phonemes present in a word. Third, we extracted syntactic features from the transcript using spaCy (Honnibal et al., 2020). These features include 45 dependency relations and 50 part-of-speech tags, resulting in a 95-dimensional binary vector. Fourth, we used spaCy’s *en_core_web_lg* model to extract 95-dimensional non-contextual word embeddings (where the same word is assigned the same embedding regardless of context). Fifth and finally, we used HuggingFace (Wolf et al., 2020) to extract LLM contextual embeddings from the activations of the middle layer of GPT-2-XL (Radford et al., 2019). These feature spaces form the regressors we used to predict the ECoG data.

We epoched the high-gamma band electrode time series per word from –2 seconds to +2 seconds (Figure 1D). Thus, the linear regression model predicts the word-by-word fluctuations in neural activity at a specific lag. We used two-fold cross-validation to train separate encoding models for each feature space, electrode, and lag on one half of the podcast stimulus. Since different feature spaces are of different dimensionality, we used ridge regression where the penalty hyperparameter is learned within the training set using the *himalaya* python library (Dupré La Tour et al., 2022). We evaluated the performance of each encoding model on the held-out fold by correlating the model-predicted neural activity with the actual neural activity across words in the test fold. Finally, we averaged the two correlations for each fold to obtain one correlation value denoting the encoding performance for each feature space, electrode, and lag.

We visualize encoding model performance both spatially (Figure 3A) and temporally (Figure 3B). We found that the acoustic and phonetic feature spaces performed well in the early auditory cortex (EAC) and mid-superior temporal gyrus (STG). Syntactic features and non-contextual embeddings performed well along the STG and extended into the inferior frontal gyrus (IFG). LLM contextual embeddings achieved the highest encoding performance and for the most number of electrodes. The different feature spaces also yielded different temporal profiles across lags relative to word onset. For example, LLM embeddings had high encoding performance across the four-second window, whereas other feature spaces peaked after word onset (Figure 3B). In particular, the acoustic and phonetic features peaked sharply after word onset. We also observed certain temporal differences in specific regions. For example, LLM embeddings peaked sharply after word onset in IFG, but more broadly in STG (Figure 3C). Moreover, acoustic features performed better than LLM embeddings in EAC, at and after word onset. Altogether, these results validate our procedures for stimulus feature extraction, ECoG preprocessing, and temporal alignment between the ECoG signals and stimulus. Our findings also suggest that this dataset can serve as a useful resource for testing different models of language processing.

**Figure 3.**
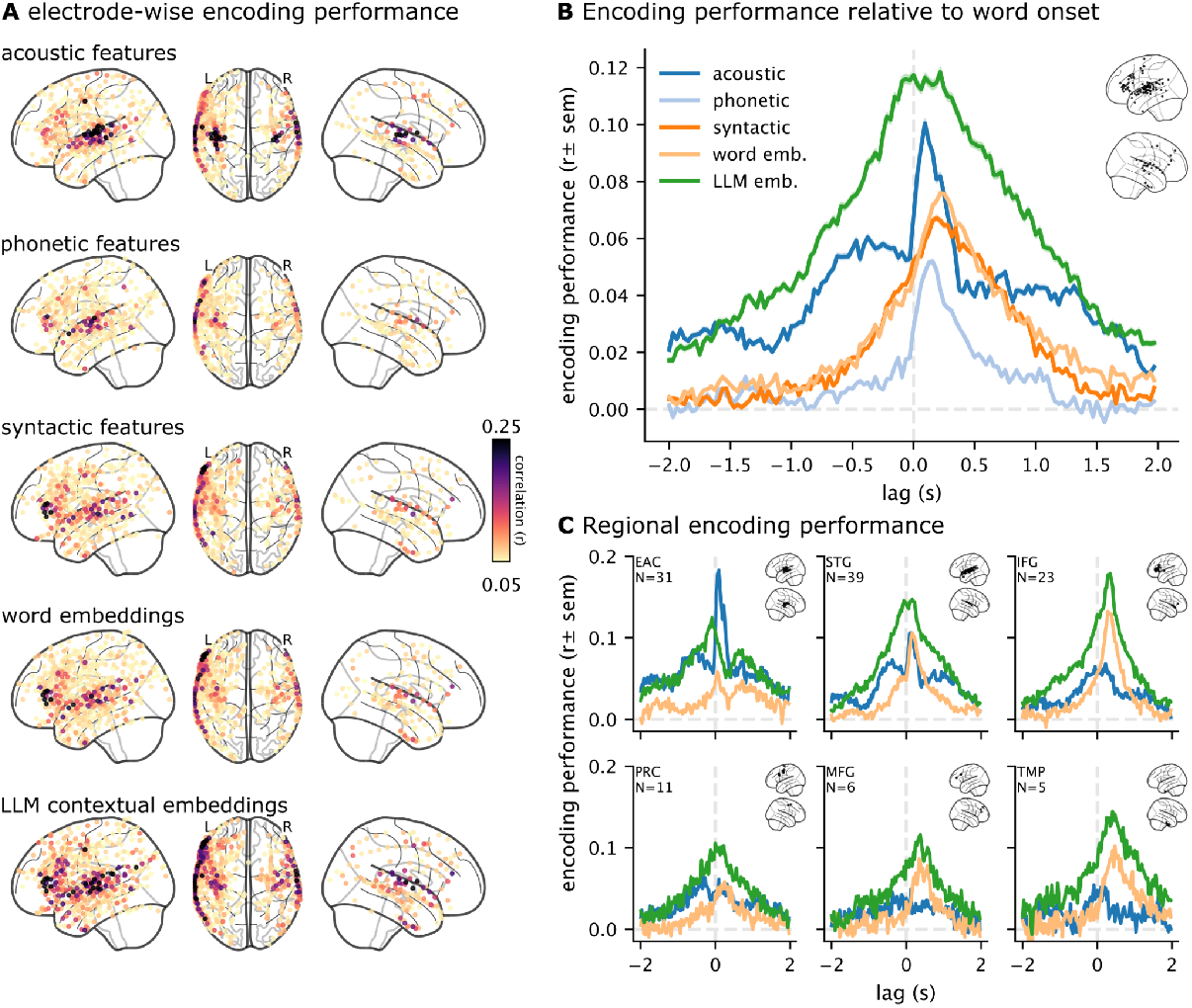
Encoding model performance for five feature spaces. (**A**) Maximum encoding performance (correlation between model-predicted and actual neural activity) across lags at each electrode for five different feature spaces. Electrodes with significant performance (*p* < .01, FDR corrected) for any of the feature spaces are visualized. (**B**) Comparing encoding performance across five different feature spaces and lags relative to word onset. The average over all selected electrodes per feature space is plotted. (**C**) Encoding performance for the acoustic, non-contextual word embeddings, and LLM embeddings are visualized for six regions of interest: early auditory cortex (EAC), superior temporal gyrus (STG), inferior frontal gyrus (IFG), precentral (PRC), middle frontal gyrus (MFG), and temporal pole (TMP). Electrodes are assigned regions based on the Destrieux atlas (Destrieux et al., 2010).

### Usage Notes

We include tutorial notebooks to serve as pedagogical examples for using the data and replicating the results presented in this paper. These tutorials are available online at https://hassonlab.github.io/podcast-ecog-tutorials and in the supplementary materials. We encourage researchers to familiarize themselves with the BIDS-iEEG (Holdgraf et al., 2019) modality-specific standards to best use the dataset. We also recommend the use of MNE tools (Gramfort, 2013) for reading the data (e.g., the preprocessed .*fif* files and to facilitate analyses). The tutorials cover the following topics: downloading data and installing the required libraries, preprocessing ECoG data, running quality control analyses, extracting stimuli features, and performing encoding analyses.

In this paper, we introduced the “Podcast” ECoG dataset and replicated several previously published analyses. That said, many of these analyses include several simplifying choices and may overlook certain questions of interest. For example, we epoch the neural activity word-by-word and train separate models at different lags. Linguistic information may be encoded in finer-grained subword dynamics or dynamics evolving over the course of word articulation. An alternative method may bypass epoching altogether and perform a time-resolved analysis similar to encoding models in fMRI (Huth et al., 2016). Furthermore, we index our analyses to the onset of words.

However, words have different lengths and internal structures; anchoring to the center of each word may provide a good compromise (Mischler et al., 2024). Models that aggregate features across words (e.g., sentence embeddings; (Muennighoff et al., 2022)) may also provide novel insights relative to our word-by-word analysis. Our analyses use high-gamma band power as an index of neural activity. Linguistic information may also be encoded in oscillations at other frequency bands or in cross-frequency coupling (e.g., Weissbart & Martin, 2024). Finally, we recommend using a variance partitioning analysis (Lescroart et al., 2015) to measure the unique contribution of a particular feature set, such as LLMs, relative to other features (see our tutorials for an example).

## Code Availability

Data and code associated with data curation are available on OpenNeuro (https://openneuro.org/datasets/ds005574). The code for analyses in this paper is available on GitHub (https://github.com/hassonlab/podcast-ecog-paper). Tutorials are also available on GitHub (https://hassonlab.github.io/podcast-ecog-tutorials).

## Acknowledgments

We thank Catherin Chen and Amir Khalilian for the discussion regarding ECoG preprocessing.

## Author contributions

Conceptualization: ZZ

Data curation: ZZ, BA

Formal analysis: ZZ

Funding acquisition: UH, AF

Project administration: SA, UH

Software: ZZ

Visualization: ZZ

Writing – original draft: ZZ

Writing – review & editing: ZZ, SA, UH

